# Modern clinical *Mycobacterium tuberculosis* strains leverage type I IFN pathway for a pro-inflammatory response in the host

**DOI:** 10.1101/594655

**Authors:** Deepthi Shankaran, Prabhakar Arumugam, Ankur Bothra, Sheetal Gandotra, Vivek Rao

**Author notes:** Equal contribution (Deepthi Shankaran, Prabhakar Arumugam).

## Abstract

Host phagocytes respond to infections by innate defense mechanisms through metabolic shuffling in order to restrict the invading pathogen. However, this very plasticity of the host provides an ideal platform for pathogen mediated manipulation. By employing the macrophage model of Mtb infection, we identify an important strategy employed by modern clinical lineages in regulating the host immune–metabolism axis. The potent ability of these strains to specifically elicit a strong and early macrophage type I IFN response (in contrast to the protracted response to ancient Mtb), was dependent on an increased ability to localize in acidified phagosomes; this higher transit via acidified compartments is important for stimulation of the DNA dependent signaling in infected macrophages. The augmented IFN signaling provided a positive regulatory loop for enhanced expression of the cellular oxysterol-CH25H which in turn facilitated higher levels of IL6 in macrophages infected with the modern Mtb strains. Requirement of type I IFN signaling in mycobacterial intracellular growth highlights another unique ability of Mtb to manipulate host cell physiology and proinflammatory responses.

**Significance Statement:** Co-evolution with humans has enabled the development of novel adaptive mechanisms for survival in host specific environments in the human TB pathogen-Mtb. We present one such mechanism of modern Mtb strains harnessing the type I IFN immune axis to regulate the host pro-inflammatory response. Our results highlight the use of host intracellular endosomal transit as a mechanism by these strains to ensure a strong type I IFN response in macrophages. We also demonstrate the ability of Mtb to regulate macrophage cholesterol metabolism in order to fine tune the host innate responses. These findings lay the foundation of the future development of a host axis directed intervention strategy against this pathogen.

## Introduction

A long-standing association with the human population has been critical to the development of specific immune modulatory mechanisms by *Mycobacterium tuberculosis* (Mtb) for efficient survival within the host. With the advent of high throughput genome analysis, Mtb strains have been differentiated into 7 distinct groups that can be classified into phylogenetically ancient (restricted geographic penetration) and modern (widespread) lineages. Analysis of strain specific differences in their virulence and disease manifestation has led to identification of possible molecular correlates of virulence (*1-3*).

The initial contact of Mtb with macrophages activates the cGAS-STING-nucleic acid signaling pathway leading to type I IFN expression (*4*, 5). This response is mediated by host factors like ROS induced mitochondrial damage and pathogen associated factors like ESX-1 mediated phagosomal escape and mycobacterial cyclic di-nucleotides (*6-10*). A detrimental pathogenic effect has been associated with Mtb induced type I IFN signaling in regulating the expression of cytokines such as TNFα, IL12 and IL10 in macrophages (*11-13*). Moreover, type I IFNs have been shown to alter the IFN γ dependent activation of macrophages and consequent mycobacterial growth (*14*). However, another study has also suggested a beneficial role for the mycobacteria induced type I IFN signaling in IL12 production and antigen presentation by macrophages (*15*).

Clinical Mtb lineages vary significantly in their inflammatory response induction and immune modulatory functions (*16-18*). Given the duality of the outcome of type I IFN induction by Mtb in macrophages, mechanisms underlying strain specific type I IFN is still unclear. We demonstrate that the higher type I IFN induction (IFNβ) by modern lineages (L3, L4 and L2) is dependent on bacterial DNA presentation to the cGAS-STING signaling pathway as a result of an early and increased bacterial trafficking via the endo-lysosomal pathway. Consistent with this augmented IFNβ and enhanced expression of the IFN stimulated, immunomodulatory (pro inflammatory) oxysterol-CH25H, significantly higher levels of IL6 were observed in macrophages infected with the modern Mtb strains. This study provides evidence for a novel mechanism of the plasticity of modern Mtb by coupling type I IFN with macrophage pro-inflammatory response. Understanding the physiological consequence of this phenomenon would provide important insights into the successful adaptation of modern Mtb to the host environment.

## Results

### Mtb lineages induce differential type I IFN response in macrophages

The type I IFN response of macrophages has generated interest as a pro-pathogenic response induced by Mtb in macrophages (*19-21*). The THP1-Dual cell line can be used to report on the induction of Type I IFN as it expresses a secreted luciferase reporter gene under the control of an ISG54 minimal promoter in conjunction with five IFN-stimulated response elements. Detectable levels of secreted luciferase as a read out for IFN activation was observed in a temporal manner in response to Mtb Erdman at a MOI of 5 with 10-15 folds activity by 24h and 48h (Fig. 1A).

**Fig. 1:**
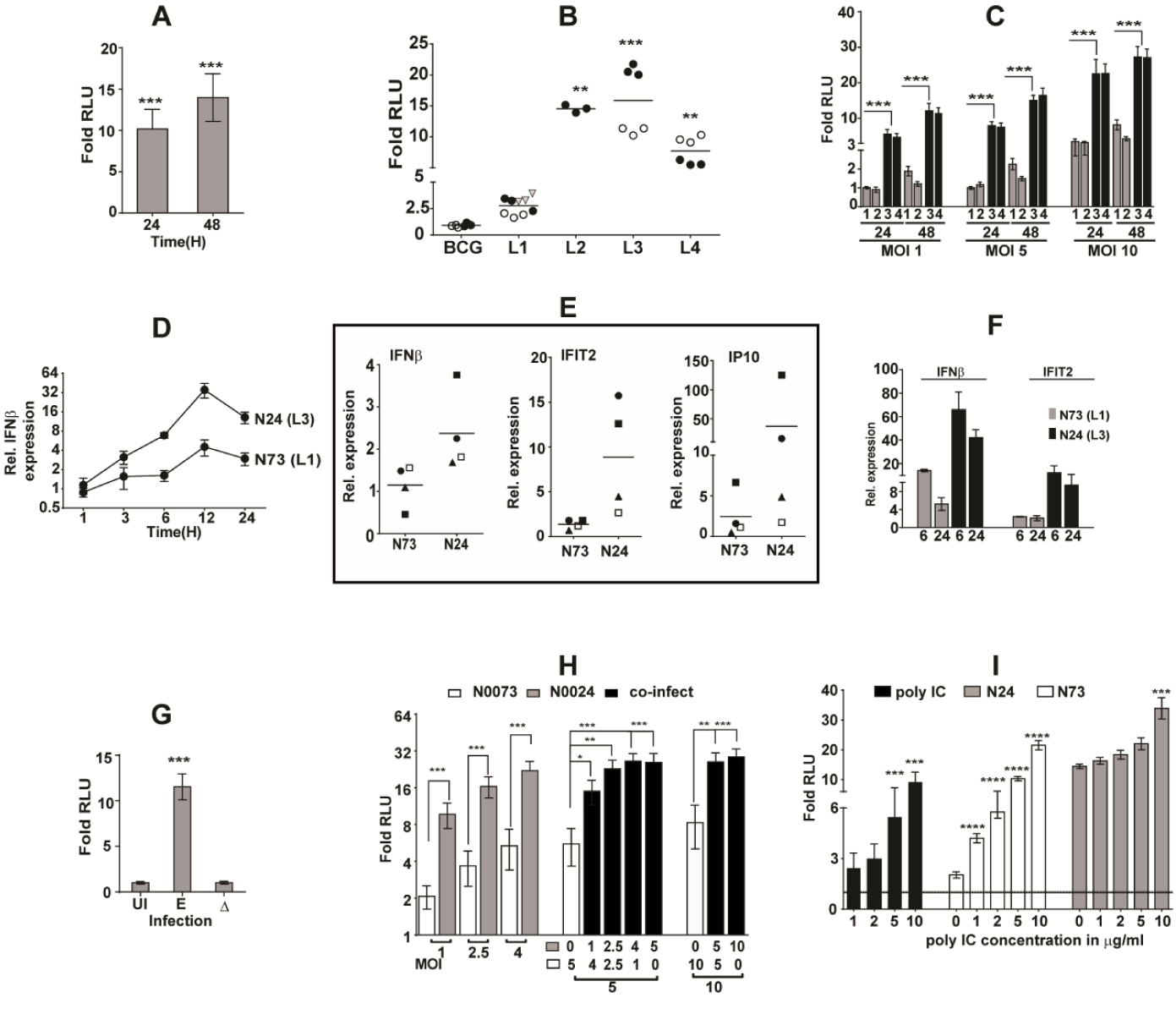
Modern lineages have evolved to induce a significantly higher type I IFN response. A-C) Relative luciferase activity in naïve and stimulated THP1 dual macrophages: Infected with Mtb Erdman at a MOI-5 (A), or with Mtb strains belonging to different lineages [L1-4] at a MOI-5 (B), 2 independent isolates of each lineage are represented as open and closed circles, T83 Vietnamese L1 subtype Mtb is shown as inverted triangle, or with Mtb L1 [grey bars; 1-N72, 2-N73] and L3 [black bars; 3-N4, 4-N24] at different MOIs (C). At the indicated time points, culture supernatants were harvested and tested for luciferase expression. Values represent average ratios ± SE of RLU in infected and control samples from triplicate wells of two independent experiments [N=2]. D-F) Relative expression in macrophages infected with N24 [L3] or N73 [L1] at a MOI-5. D) IFNβ expression at different time intervals post infection, E) IFNβ, IFIT2 and IP10 mRNA in primary MDMs from four separate individuals (symbols) and F) expression of IFNβ and IFIT2 mRNA in RAW macrophages. The average relative values ± SE (with respect to GAPDH) in infected and control samples from triplicate wells of multiple replicate experiments are represented. G-I) Relative luciferase activity in naïve and stimulated THP1 dual macrophages: Infected with live or heat killed (Δ) Mtb Erdman at MOI-5 (G) or with N24, N73 or in different combinations (H) or stimulated with poly IC alone or in combination with Mtb at a MOI of 5 (I).

Further, we also observed a distinct pattern of response in these macrophages infected with clinical Mtb strains of different phylogenetic lineages. Macrophages infected with the modern Mtb lineages (L2-HN878, L3-N4, N24; L4-H_37_Rv, Erdman, H_37_Ra) displayed 8-20 folds higher luminescence activity at 24h of infection contrasting with the lower (∼2.5 folds) or complete absence of activity in macrophages infected with the ancient lineage I (T83, N72, N73) Mtb and BCG strains, respectively (Fig. 1B, Fig. S1). This higher activation potential of L3 Mtb was maintained across different infective doses (Fig. 1C). While L3 Mtb strains (N4 and N24) could activate this response even at a low MOI of 1 (4.6 folds) by 24h of infection, these levels were higher than the average levels of IFN response induction (∼3 folds) by L1 Mtb (N72 and N73) at MOI-10.

Gene expression profiles further validated the temporal and differential IFNβ expression profiles of Mtb infected macrophages, with the L3 Mtb strain-N24 beginning to induce IFNβ expression as early as 1h with a gradual increase (∼30 folds by 12h of infection) that was sustained even by 24h of infection (∼16 folds) (Fig. 1D). In contrast, the L1 strain (N73) induced IFNβ at a significantly later time point in infection (6-12h) and at 5-7-fold lower levels even by 24h of infection.

This pattern of an attenuated IFN response by L1 Mtb was also evident in primary human MDMs and the mouse macrophage line RAW264.7. Consistently, L3 Mtb induced significantly higher (4-10 folds) IFN response than L1 Mtb at any point in infection (Fig. 1E, F). The induced IFN response required active infection as macrophages infected with non-viable Mtb failed to initiate this response (Fig. 1G). To verify if L1 Mtb actively suppressed type I IFN response, we employed a co-infection assay of N24 and N73 in different ratios (Fig.1H). At all doses, N24 induced significantly higher response than N73, however, in mixed infections, presence of four-fold excess N73 (N73:N24::4:1) could not diminish the response induced by N24. Consequently, IFN levels in mixed infections were comparable to levels in macrophages infected with N24 alone, even at the overall higher MOI of infection of 10. Subsequently, in a trans-activation/ suppression assay, again N73, did not suppress poly IC mediated IFN response (Fig. 1I). In most cases, response to poly IC was clearly augmented in conditions of co-stimulation with either N24 or by N73.

### IFN inducing capability correlates with induction of a pro-inflammatory response via expression of CH25H in macrophages

To test if L1 Mtb strains were attenuated in overall macrophage activation, we tested gene expression and cytokine profiles of RAW264.7 macrophages infected with L1 and L3 Mtb. Both TNFα and IL6 patterns were strikingly similar to type I IFN, with lower levels in the case of L1 than the L3 infected RAW264.7 macrophages after 24 hours (Fig. 2A). This attenuated IL6 and TNF expression was also observed in bone marrow-derived macrophages from C57BL/6 mice with L1 Mtb inducing minimal amounts of the cytokines (Fig. 2B). L3 Mtb induced significantly higher levels of IL6 in a temporal manner attaining peak values by 48h of infection as opposed to minimal levels of cytokines even by 72h of L1 Mtb infection (Fig. 2C). A similar pattern of low IL6 induction by L1 Mtb in human monocyte derived macrophages of 4 healthy individuals at 6h and 24h of infection suggested a universal mechanism of regulation in the macrophage response pathway by this mycobacterial strain (Fig. 2D).

**Fig. 2:**
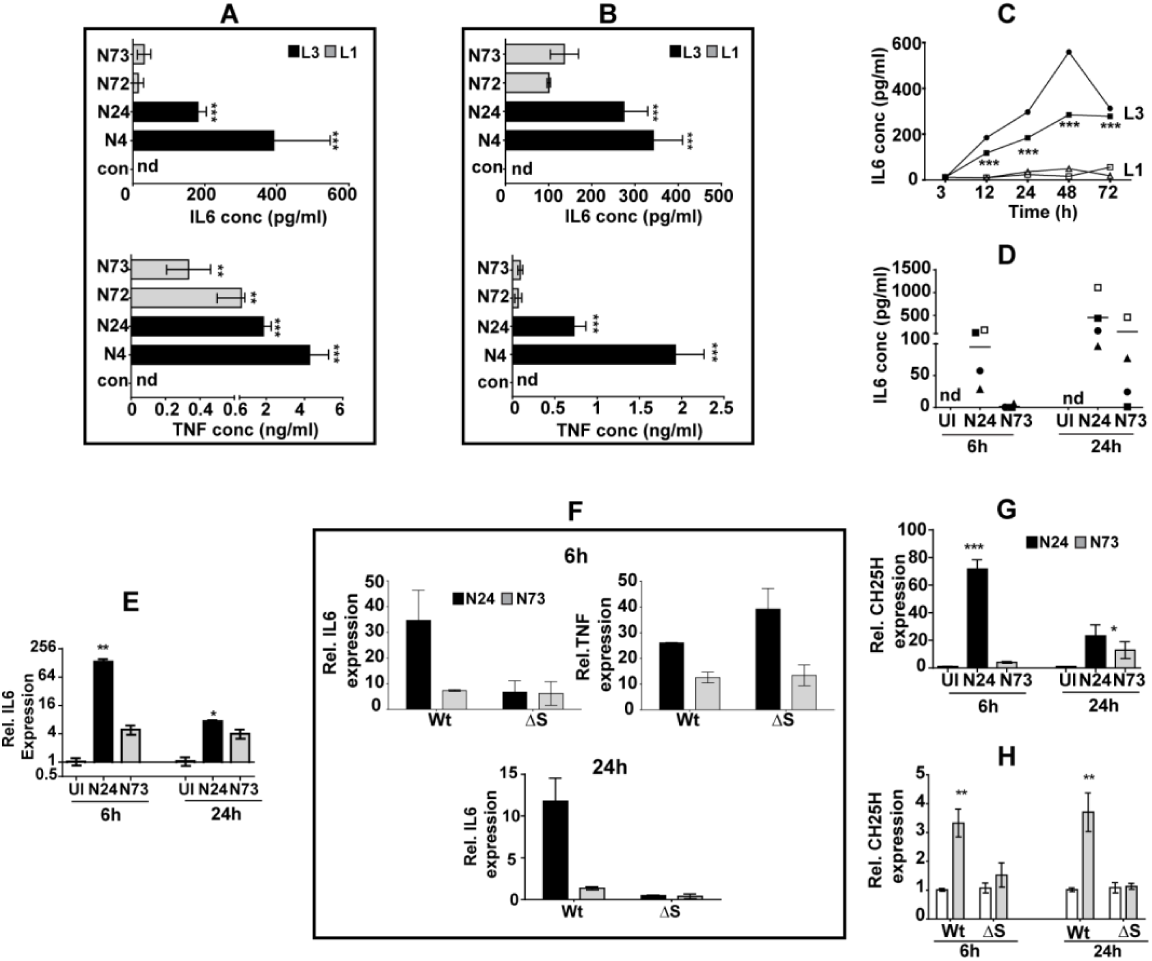
IL6 response in Mtb infected macrophages primarily depends on STING mediated CH25H expression. A-C) Levels of cytokine secreted from RAW264.7 macrophages (A), BMDMs at 24h (B) or at different time intervals (C) following infection with two L3 or L1 Mtb strains. Values represent mean cytokine concentration in pg/ml ± SEM of triplicate assays (N=3). D) Levels of IL6 in Mtb infected MDMs from four individuals at 6h and 24h *p.i*. Values represent mean cytokine concentration in pg/ml in the culture supernatant of triplicate wells for each individual. E, F) IL6 and TNF gene expression profiles in BMDMs (E) and RAW264.7 macrophages (F) following infection with N24 (L3) or N73 (L1) Mtb strains. The average relative IL6 values ± SE (with respect to GAPDH) at 6h and 24h post infection in triplicate assays is shown. G, H) Expression of CH25H in BMDMs (G) and RAW macrophages (H) following infection with Mtb strains at 6h and 24h *p.i*. at MOI-5. The average relative IL6 values ± SE (with respect to GAPDH) at 6h and 24h *p.i*. in triplicate assays of (N=2/3) is shown.

To check if the regulation was at the transcript level, we compared gene expression by qPCR. In macrophages infected with N24, ∼30 fold higher *il6* expression was observed than in N73 infected cells as early as 6h *p.i*. While the expression declined at a later time point (24h) in these macrophages, the overall levels were significantly higher than in the N73 infected cells (Fig. 2E).

Both strains were capable of inducing *il6* by 6h; N24 by 34 folds and N73 by 7 folds. While we did not observe any change in the levels of *il6* expression in the N73 infected ΔSTING macrophages, the differential higher response (∼5x) seen in the case of N24 infection was lost (Fig. 2F). Although, *il6* levels were lower at 24h of infection with either Mtb strain, loss of STING abrogated expression in macrophages suggestive of an inherent regulation of IL6 by IFN1 in macrophages. More importantly, loss of STING did not significantly affect the TNF inducing capacity of Mtb in macrophages. Recent evidence points to a positive regulatory circuit for IL6 via cGAS-STING regulated expression of CH25H, responsible for the synthesis of the pro inflammatory oxysterol, 25 hydroxy-cholesterol (*22-23*). Examination of genes involved in cholesterol biosynthesis revealed a temporal but distinct pattern of expression in L3 and L1 Mtb infected BMDM (Fig. S2). Most of the genes involved in the cholesterol biosynthetic pathway were either up-regulated or similar in L1 infected macrophages after 24h of infection. In contrast, these genes were inhibited in response to infection with L3 Mtb. Surprisingly, in contrast with the overall pathway, the expression of CH25H was significantly upregulated in the L3 infected macrophages (60-70-fold higher than the uninfected control) as opposed to a much lower expression of this gene (3-4 folds) in the L1 infected macrophages, by 6h of infection (Fig. 2G). This difference was, however, lower at 24h, wherein a sharp decline in L3 (23 folds) and an increase in L1 infected macrophages (10-12 folds) was observed. Given the increased CH25H expression and requirement of type I IFN for enhanced IL6 expression, we tested if STING signaling contributed to the early increase of CH25H in L3 Mtb infected macrophages. The complete loss of CH25H expression in ΔSTING macrophages substantiates our hypothesis of a regulatory role for type I IFN in host oxysterol mediated IL6 response in modern Mtb infected macrophages (Fig. 2H).

### L1 Mtb are delayed in inducing type I IFN response in macrophages despite the presence of an active ligand

Previous studies have implicated an important role of the Mtb ESX-1 secretion system in cytosolic egress of ligands and activation of the cytosolic surveillance pathway resulting in type I IFN activation (*5-10*). However, we did not observe any difference between the IFN inducing abilities of H_37_Ra or H_37_Rv despite the reported differences in the levels of the Esx-1 proteins (*24*). Further, comparable expression of ESAT6 in our clinical Mtb strains (L1 and L3) (Fig. S3A) argued for an esx-1 independent mechanism of Type 1 IFN induction.

We also did not observe any difference in mitoSOX staining of macrophages infected with either Mtb strain, contrasting with the previously identified mode of mitochondrial ROS mediated type I IFN signaling (*10*) hinting at alternative early events associated with recognition of bacterial ligands (Fig. S3B).

Having ascertained that N73 did not suppress IFN induction, it was logical to assume lack/ lower quantities of the active ligand as an underlying cause for the deferred IFN response by L1 Mtb. In accordance with previous reports, Mtb induced type I IFN response of macrophages was completely dependent on DNA mediated signaling. Wt RAW-luc (ISG) cells also portrayed a differential type I IFN response to Mtb lineages, with 5-10-fold higher IFN by L3 Mtb in comparison to L1 Mtb strains. Only the loss of cGAS, STING and TRIF (∼60% decrease), associated with DNA mediated signaling as opposed to RNA sensing, diminished the type I IFN response to L3 Mtb (Fig. 3A, S4A). Lipofectamine based transfection of DNA results in delivery of DNA to the endosomal system, and eventually to the cytosol via formation of multiple transient pores in the endosomal membrane (*25*). As cytosolic sensors such as STING are essential for eliciting this response in case of Mtb infection (*10, 26, 27*), we argued that delivery of mycobacterial DNA by lipofectamine may also be able to elicit this response. In fact, with as low as 125 ng of genomic DNA from Mtb, macrophages responded with IFN expression that was again abolished in cells stimulated with DNAse treated DNA (Fig. 3B). Surprisingly, genomic DNAs from N73 as well as BCG were similar to Erdman and N24 in their IFN inducing capacities (Fig. 3C). Given the presence of an active molecule (DNA) even in Mtb strains with attenuated IFN induction, it was reasonable to assume the lack of DNA presentation as the basis for lower IFN. A preliminary evaluation hinted at a temporal IFN response to Mtb Erdman extract (cell lysates) when delivered to THP1 cells via transfection, with the response increasing from ∼10 folds at 3h to ∼75 folds by 6h and then declining to ∼10-fold expression by 24h of stimulation with different doses (Fig. 3D). Even though very low amounts of the extract (1.6 ng) was sufficient to induce ten-fold IFN by 6h (Fig. S4B), the complete loss of IFN only in extracts subjected to DNAse digestion argued for the pivotal role of DNA sensing in Mtb induced type I IFN in macrophages and its inaccessibility to the IFN signaling machinery of macrophages (Fig. 3E). Further proof of effective ligand masking by L1 Mtb and BCG came from our observation that delivery of BCG with lipofectamine was sufficient to enhance the Type I IFN response by 6 folds compared to BCG infection without lipofectamine (Fig. 3F).

**Fig. 3:**
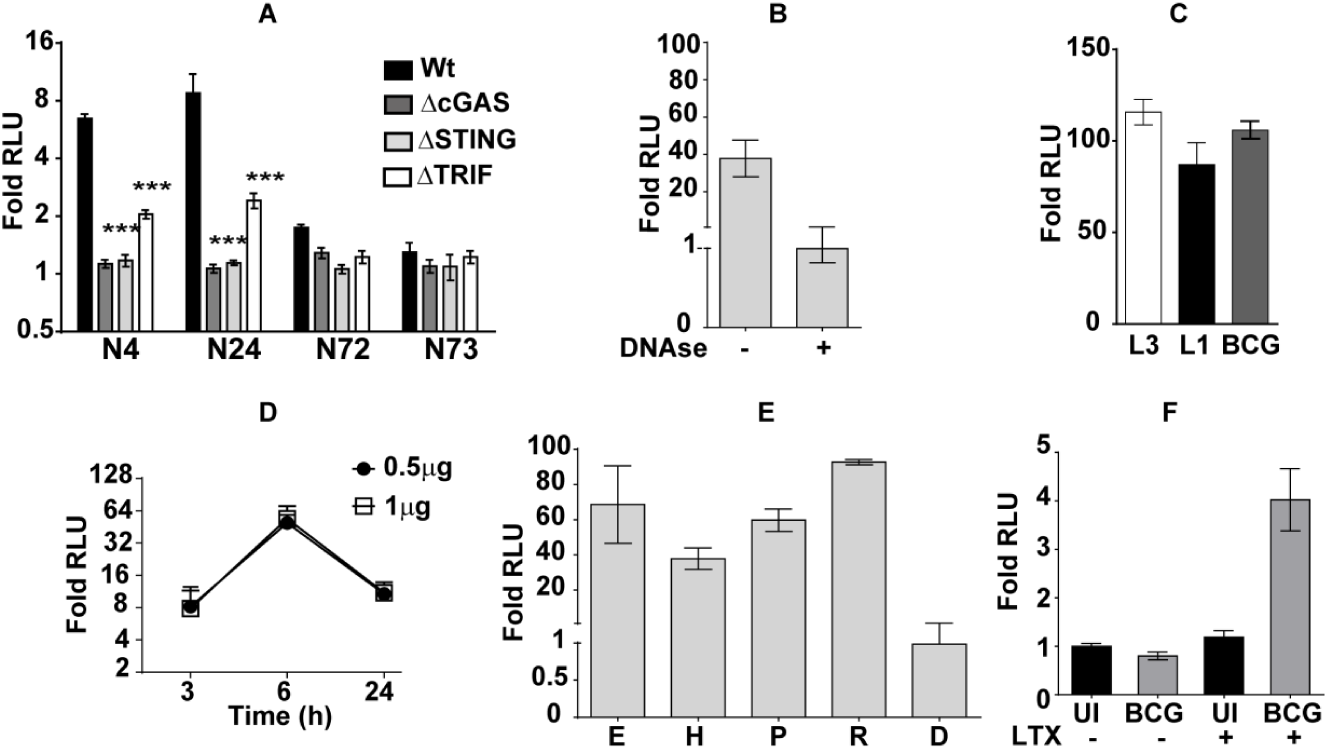
Strains with suppressed type I IFN response are well equipped with the active IFN inducing ligand. A) Relative luciferase activity in Wt-ISG or mutant RAW macrophages uninfected or infected with Mtb strains belonging to L3 / L1 lineages at a MOI of 5 for 24h. Values represent average ratios ± SE of RLU in infected and control culture supernatants from triplicate wells of three independent experiments [N=3]. B-F) Relative luciferase activity in naïve and stimulated THP1 dual macrophages: following transfection with genomic DNA (B), or genomic DNA of mycobacteria (C) or crude Mtb Erdman crude lysate (D, E), or BCG transfection (F). H, P, R, D-represent treatment of extract with Heat, proteinase K, RNase and DNase, respectively. Values represent average ratios ± SE of RLU in infected and control culture supernatants from triplicate wells of multiple experiments [N=2, 3].

### Early localization to acidified compartments is important for Mtb induced type I IFN induction

The early induction of type I IFN in L3 infected macrophages advocated for molecular events that ensue immediately after infection as causative. Further, analysis of THP1 macrophages by TEM revealed the differential localization of the Mtb clinical isolates in distinct intracellular compartments-N24 localized to compact double membranous vesicles at 24h in contrast to N73 that were confined in larger less defined vesicular structures with intracellular particles (Fig. 4A). In support of this observation, we observed that Mtb N24 localized to lysotracker positive (lyso+) vesicles at significantly higher numbers than N73 at 24h *p.i*. (Fig. 4B). This differential localization to lyso+ vesicles between N24 and N73 was also observed in RAW264.7 macrophages. In a temporal analysis of Mtb transit via the endosomal pathway, differential localization between Mtb strains arose as early as 1h of infection with more than two-fold N24 in the lyso+ vesicles as compared to N73 (Fig. 4C). Despite a gradual decrease of N24 in lyso+ positive vesicles with time, consistently 2-fold higher numbers of N24 was seen throughout the experiment duration.

**Fig. 4:**
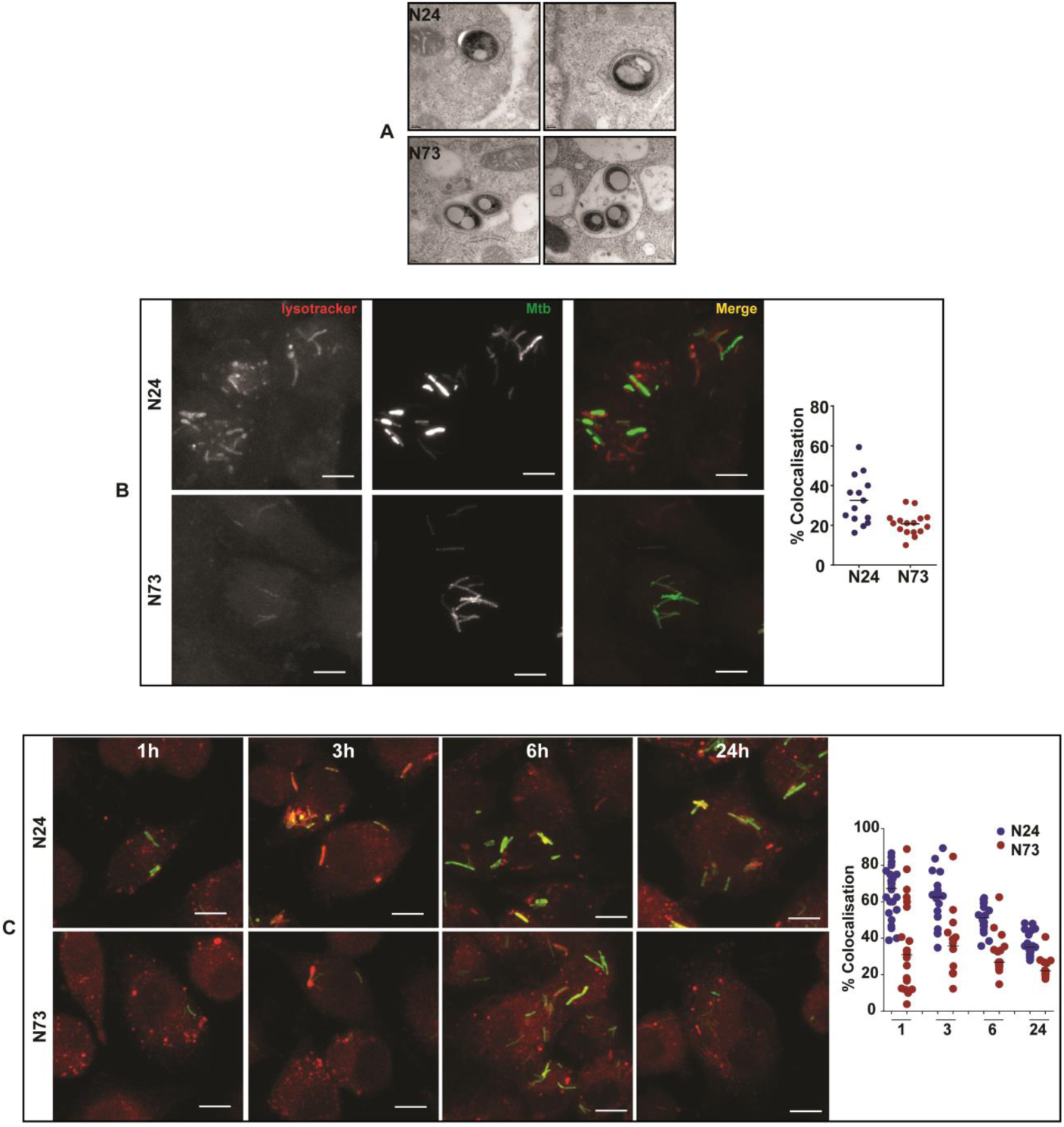
Early and sustained localization of the high IFN inducing Mtb strains with acidified vesicles. A) Representative TEM images (2) of Mtb N24 and N73 in THP1 macrophages at 24h post infection. While N24 is found in compact vesicles, N73 localizes in enlarged vesicles (black arrow) with cellular debris. B-E) Localization of Mtb N24 and N73 in acidified vesicles. Representative images of macrophages stained with lysotracker red and infected with GFP expressing N24/ N73 - THP1 (B) at 24h or RAW 264. 7 (C) at 1h, 3h, 6hand 24 h post infection is depicted. The extent of co-localization of Mtb with the lyso+ vesicles in THP1, 24h and RAW264.7 at 1h, 3h, 6h and 24h is represented graphically. Each dot represents a field of image from 2 independent experiments (> 100 infected cells).

We have previously demonstrated that L1 strains harbor a mutation in papA2, rendering them incapable of sulfolipid synthesis (*28*). A dominant role for the Mtb sulfolipids has also been demonstrated in modulation of the cellular phagosome maturation and acidification (*29-31*). We questioned whether the lack of sulfolipids renders Mtb incapable of inducing the type I IFN response. To test this hypothesis, we deleted *mmpL8*, coding for an exporter of mature sulfolipids to the bacterial cell surface and measured induction of the Type I IFN response in the reporter cells. There was no loss of IFN inducing ability of Mtb in the absence of *mmpL8* (Fig. S5), supporting an alternative reason for the differential induction of the type I IFN response. Another finding supporting the lack of sulfolipid not to be responsible for this phenotype is the inability of the Vietnamese strain of L1 lineage to induce significant Type I IFN response despite expressing sulfolipids in the membrane (Fig. 1B).

### IFN inducing capacity of Mtb relies on efficient acidification of the phagosome

Given the high numbers of N24 localization to lyso^+^ compartments, and comparable intracellular growth rates of N24 and N73 (data not shown), it was reasonable to assume an enhanced propensity of N24 to be better equipped for survival in this subcellular niche. The ability of Mtb to counter phagosomal-lysosomal fusion and acidification events is well recognized (*32-33*). To understand if phagosomal acidification was important for type I IFN induction, we treated macrophages with an established phagocytosis activation signal (LPS+ IFNγ) or phagosomal acidification blockers-bafilomycin A and chloroquine prior to infection with Mtb. Activation of macrophages with IFNγ and LPS enhanced the presence of Mtb in lyso^+^ vesicles (Fig. 5A). Importantly, the numbers of lyso+ N73 Mtb increased to comparable levels as N24 Mtb by 6h of infection (Fig. 5B). Consequently, a 10-fold higher type I IFN response was observed in N73 infected IFNγ+ LPS pretreated macrophages leading to loss of the differential IFN response between strains (Fig. 5C and inset panel). While, addition of IFNγ and LPS significantly boosted type I IFN induction even in N24 infected macrophages, treatment with phagosome acidification blockers like bafilomycin A or chloroquine on the contrary, stunted (∼5 and 2 folds, respectively) the IFN response in these cells (Fig. 5D) emphasizing the importance of phagosomal trafficking and acidification in Mtb mediated IFN induction in macrophages.

**Fig. 5:**
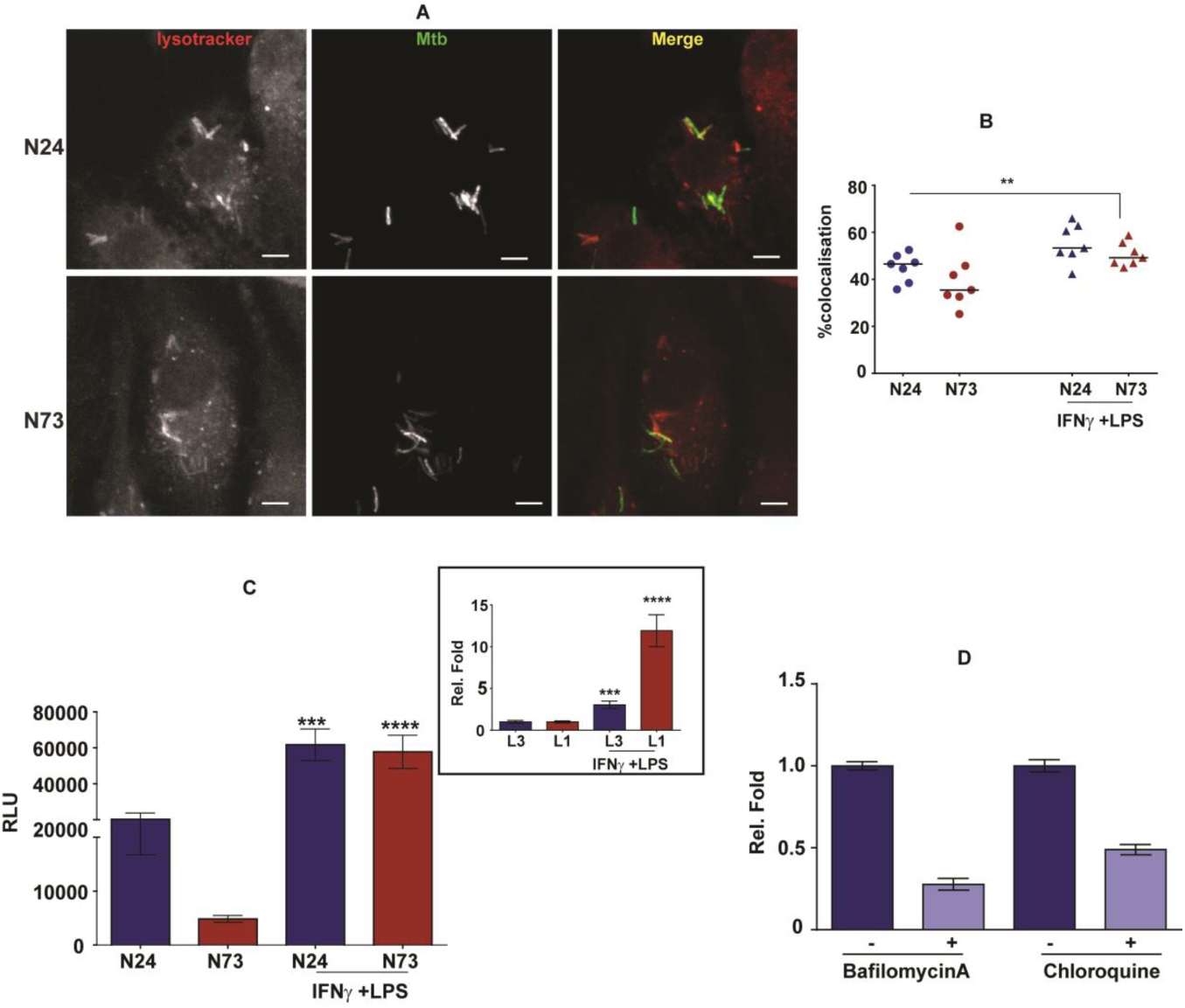

### Abrogation of DNA signaling arrests mycobacterial growth in macrophages

In an effort to test the physiological relevance of type I IFN, we analyzed the ability of Mtb N24 to grow in Wt ISG and macrophages deficient in STING or TRIF. Over 3 days of culture, Wt macrophages supported Mtb growth (Fig. 6). In contrast, despite, similar levels of uptake in the deficient macrophages, Mtb N24 growth was restricted to initial levels at all times (bacteriostasis) strongly arguing for an important role for the early type I IFN signaling in intracellular growth of Mtb.

### The working model

Clinical Mtb strains differ in their intracellular trafficking routes to arrive at or avoid acidified compartments; modern lineages traffic to arrive early into acidified compartments of host macrophages. The reduced movement of ancient Mtb strains likely hinders the access of Mtb DNA to cytosolic DNA sensors, thereby limiting the nucleic acid response pathway and leading to weaker type I IFN response. The higher activation of type I IFN by the modern strains manifests as a substantial upregulation of the pro-inflammatory cytokine IL6 via induction of cholesterol 25-hydroxylase in a positive feedforward loop and is essential for bacterial growth in macrophages (Fig. 6).

**Fig. 6:**
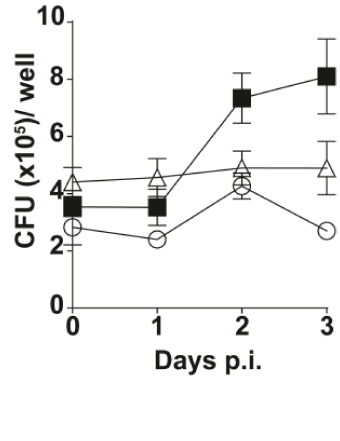
Growth of Mtb N24 in RAW reporter macrophages. Macrophages were infected with Mtb at an MOI of 5 for 6h and the intracellular bacterial numbers were enumerated by dilution plating. Average bacterial numbers / well ± SEM in WT (ISG-solid squares) or ΔSTING (open triangle) or ΔTRIF (open circle) are represented graphically at 0, 1, 2 and 3 days *p.i*.

**Fig. 7:**
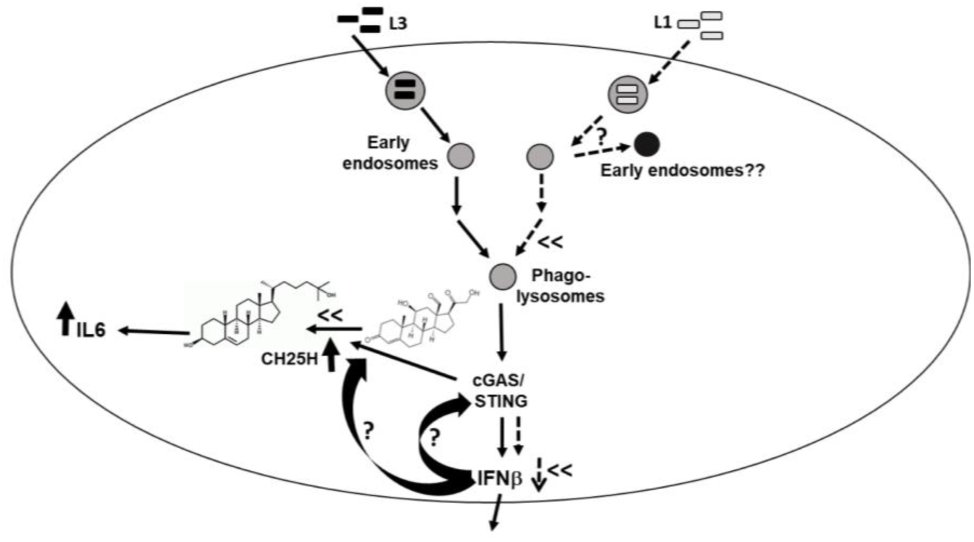
A putative working model of modern Mtb induced IFN and subsequent IL6 expression via the STING-CH25H pathway.

## DISCUSSION

The grand success of Mtb as a human pathogen stems from its extreme plasticity to suit the host cell niche, modulate the host response and survive multiple intracellular stresses. The immediate response of host phagocytes upon contact is the activation of inflammatory responses aimed at controlling the growth of the invading pathogen. Typically the type I IFN response is directed against viral infections, but recent studies have highlighted the macrophage expression of type I IFNβ in response to intracellular (pathogenic) and extracellular bacterial (protective) infections (*34-39*). We observed that modern Mtb strains are proficient in inducing this response than the evolutionarily older strains of Mtb. We also observed a strong correlation between the STING mediated IFN induction and high levels of IL6 response in macrophages, arguing for a novel mechanism of macrophage response regulation by the modern Mtb strains. Recent evidence also connects the type I IFN induced cholesterol 25-hydroxylase (CH25H) with the expression of pro-inflammatory cytokines like IL6 in macrophages (*22, 23, 40-42*). Given the lower expression of genes for cholesterol biosynthesis in the late stage of infection with N24 Mtb in line with the earlier report (*43*), it was surprising that expression levels of CH25H were on the contrary increased, signifying its importance in bridging type I IFN and IL6 expression in macrophages. Moreover, the complete loss of CH25H expression in the STING deficient macrophages following N24 infection highlights this novel regulatory loop of type I IFN-CH25H-IL6 regulation in modern Mtb infections. Overall, we define a novel pathway of pro-inflammatory cytokine expression in Mtb induced macrophages involving the early innate type I IFN response and active modulation of the host cell oxysterol metabolism. Our results identify a novel link between known pathogenicity of modern lineages (*16*) and their ability to mount a feedforward pro-inflammatory response via the Type I IFN axis. As demonstrated previously, loss of STING/ cGAS completely abrogated Mtb type I IFN response in macrophages implicating cellular DNA signaling as a predominant mechanism of this response. Given the early induction of type I IFN and the absence of this response despite the presence of active ligand, we argued that a novel mechanism that actively concealed the DNA from the cellular nucleic acid signaling mechanism was responsible for the lower IFN induction by the attenuated ancient Mtb strains. Interestingly, loss of TRIF also resulted in absence of IFN response in N24 infected macrophages suggesting a role for RNA signaling in this response. Surprisingly, we did not observe loss of IFN induction by cellular extracts treated with RNAse. Recent reports have identified a TRIF dependent activation of STING in the DNA induced IFN response of macrophages (*44*); our results demonstrate a dominant role for this signaling pathway in Mtb mediated type I IFN response.

Two separate observations supported our hypothesis that intracellular localization early in infection would be primarily accountable for this differential IFN signaling by Mtb strains: Induction of IFN response by 1) transfected BCG as opposed to direct infection and 2) absolute requirement of DNA to be transfected. Bacterial lysis by the phagosome-lysosome pathway has been implicated in several of the innate immune response pathways in phagocytes (*45-46*). Our results provide insights into the importance of transit of Mtb strains via an acidified compartment as an important determinant of IFN induction in macrophages. Based on the presence of a double membrane compartment, we speculate this to be an autophagosome like compartment. While on the one hand only live bacteria were able to stimulate the macrophage type I IFN response, dead Mtb despite their well-documented higher rates of trafficking to acidified P-L compartments were unable to induce this response in macrophages, pointing at the ability of modern Mtb strains to utilize its early localization to an acidified compartment for active presentation of DNA (in contrast to dead bacteria that fail in this presentation) to the cytosolic signaling complex. Additionally, the loss of Mtb mediated IFN response in Baf A/ chloroquine treated macrophages and a significant increase by LPS+IFNγ pretreatment reveals an important role of phagosomal acidification in type I IFN induction.

While sulfolipids have been demonstrated in promoting phagosomal acidification and L1 strains lack sulfolipids, our results do not purport an active role for this lipid in the type I IFN response. Despite the presence of mature sulfolipids in the L1 subtype Mtb strains T83 (*28*), induction of type I IFN was relatively low in THP1 macrophages. Moreover, while differences in sulfolipid expression between H_37_Ra and H_37_Rv is well established (*47-48*), there was no discernable alteration of IFN expression between the two Mtb strains. The complete lack of change in IFN induction ability of a sulfolipid deficient Erdman Mtb (ΔM8) further absolves the critical role of sulfolipid in type I IFN induction by Mtb (Fig. S5).

In summary, we identify a novel means for modern strains of Mtb to mount a pathogenic immune response via their early passage through an acidified endosomal compartment. This passage allows presentation of their DNA to cytosolic sensors, which then triggers a cholesterol driven feed forward loop of pro-inflammatory responses. An understanding of these early trafficking events in the future will help identify means to counter this pathogenic immune response triggered by the modern Mtb strains.

## Material and Methods

### Bacterial Strains and Growth Conditions

Mycobacterial and *E. coli* strains were cultured as per standard protocols with the appropriate supplements and selection antibiotics as required. The M6 promoter fragment was amplified as described earlier (*49*) and cloned upstream of GFP in the mycobacterial expression vector-pMV261 and electroplated into mycobacteria. Expression of ESAT6 in the culture filtrate and secreted Mtb cultures was performed by immunoblotting with anti ESAT6 monoclonal antibody HYB76-8 as described earlier (*3*). The *mmpL*8 null mutant (δM8) was created by homologous recombination using specialized transduction and recombineering as described earlier (*50*). Transducing phages generate by recombineering mediated generation of phages were used for transduction of Wt Erdman Mtb cultures and hyg^R^ colonies were screened for deletion by qPCR and checking expression of sulfolipid in apolar lipid fractions by TLC as per recommended protocols (*51*).

### Macrophage culture and maintenance

All the cell lines used were tested for mycoplasma contamination at regular intervals and cultured according to the manufacturer’s recommendations. THP1 cells were grown in HiglutaXL RPMI-1640 (Himedia laboratories, India) with 10% FBS and differentiated by using PMA as described earlier (*50*). Human monocyte derived macrophages (MDMs) were isolated from 20ml as described earlier (*49*). For primary mouse macrophages, bone marrow cells were harvested from the hind limbs of C57Bl6 mice and differentiated into macrophages in 20% L-cell supernatant for 7 days as described earlier (*52*).

### Preparation of Mtb fractions for stimulation of macrophages

Mtb crude extracts were prepared from logarithmic liquid cultures with similar bacterial densities for the different strains. Cells were lysed with a mixture of 0.1 µm and 0.7 µm zirconia beads in PBS using a bead beater (Biospec India). The resultant lysates were cleared off the cell debris by high speed centrifugation and sterilized by filtration and used to transfect THP1 dual cells using Lipofectamine LTX (Invitrogen, USA) reagent according to the recommended protocol. Genomic DNA was isolated according to the manufacturer’s recommendations (Nucleopore gDNA kit) and used for transfection. Enzymatic treatment of lysates was performed as: 1) Proteinase K-at 55°C for 1h, 2) DNase, RNase at 37°C for 1h and thermal denaturation at 96°C for 30min.

### Infection of macrophages with Mtb

Macrophage cultures were seeded at the requisite density and infected with single cell suspensions (SCS) as described earlier (*50*). At different time intervals, the cells/ RNA/ supernatants were harvested and used for analysis. Bacterial numbers were counted from individual wells by standard dilution plating of the lysate. Macrophages infected with fluorescent Mtb strains were used for lysosomal localization by using 100nm Lysotracker–Red DND-99 (Invitrogen, USA) according to manufacturer’s recommendations. Macrophages transfected with mito-DsRed (kind gift from Dr. Sowmya Sinha Roy) was used for analysis of mitochondrial architecture. The cells were fixed in 4% paraformaldehyde for 30 min, mounted on slides with Prolong diamond anti fade (Invitrogen, USA) and subjected to microscopy in the Leica TCS SP8 Confocal Microscope (Leica Microsystems, USA). For estimating mitochondrial ROS by Mitosox staining, macrophages were washed thrice with PBS and stained with 5µM Mitosox (Invitrogen, USA) in HBSS for 30 min at 37°C, fixed and subjected to high content microscopy in a IN Cell Analyzer 6000(GE healthcare life sciences, USA).

### Transmission Electron Microscopy

THP1 cells were infected with mycobacteria at a MOI of 5 for 24h, fixed in 2.5% gluteraldehyde and 4% paraformaldehyde, dehydrated in graded series of alcohol and embedded in Epon 812 resin. Ultrathin sections were cut and stained with uranyl acetate and lead citrate and imaged by using Tecnai G2 20 twin (FEI) transmission electron microscope (FEI, Thermo Fisher Scientific Inc, USA).

### Statistical Analysis

All experiments were performed in multiple biological replicates (N=2 or 3). Statistical analysis was done by using Student’s t-test and corresponds to a *, **, *** for p-values<0.05, <0.01and <0.001, respectively.

### Ethics Statement

The protocol for isolation of human blood PBMCs was approved by the Institutional Human ethics committee (Ref no: CSIR-IGIB/IHEC/2017-18 Dt. 08.02.2018). Blood was collected from people with their informed consent as per the Institutional guidelines.

## Abbreviations

IFN: Interferon
BAF A: bafilomycin A
MDM: monocyte derived macrophages
Mtb: *Mycobacterium tuberculosis*
BCG: *Bacille Calmette Guerin*
CH25H: Cholesterol 25-Hydroxylase
lyso^+^: lysotracker positive, L1, L3-Lineage 1,3

## Acknowledgements

The authors thank CSIR (VR-BSC0123), and DBT (GAP0096) funding agency for supporting the study. The authors thank Mr. Manish Kumar and CSIR-BSC0403 for the confocal microscopy facility. The Transmission Electron microscopy (TEM) and mass spectrometry facilities are duly acknowledged. The institute BSL3 facility project (STS0016) is duly acknowledged. The student fellowships from CSIR and DBT India are acknowledged. The support provided by Academy of Scientific and Innovative research (AcSIR) is duly acknowledged.

## Author Contributions

DS, PA, AB, SG, VR were involved in conceptualizing and design of the work, the work was performed by DS, PA and AB. VR, PA and DS were involved in the manuscript preparation and proof reading.

## Supplementary figures

**Fig S1:**
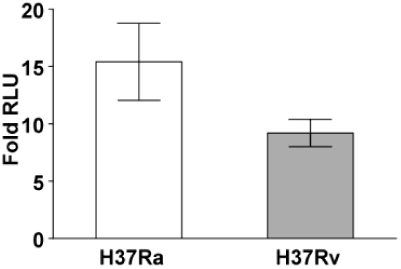
Relative luminescence in THP1 macrophages infected with H_37_Ra and H_37_Rv at 24h *p.i.* Values represent average ratios ± SE of RLU in infected and control culture supernatants from triplicate wells of two independent experiments [N=2].

**Fig S2:**
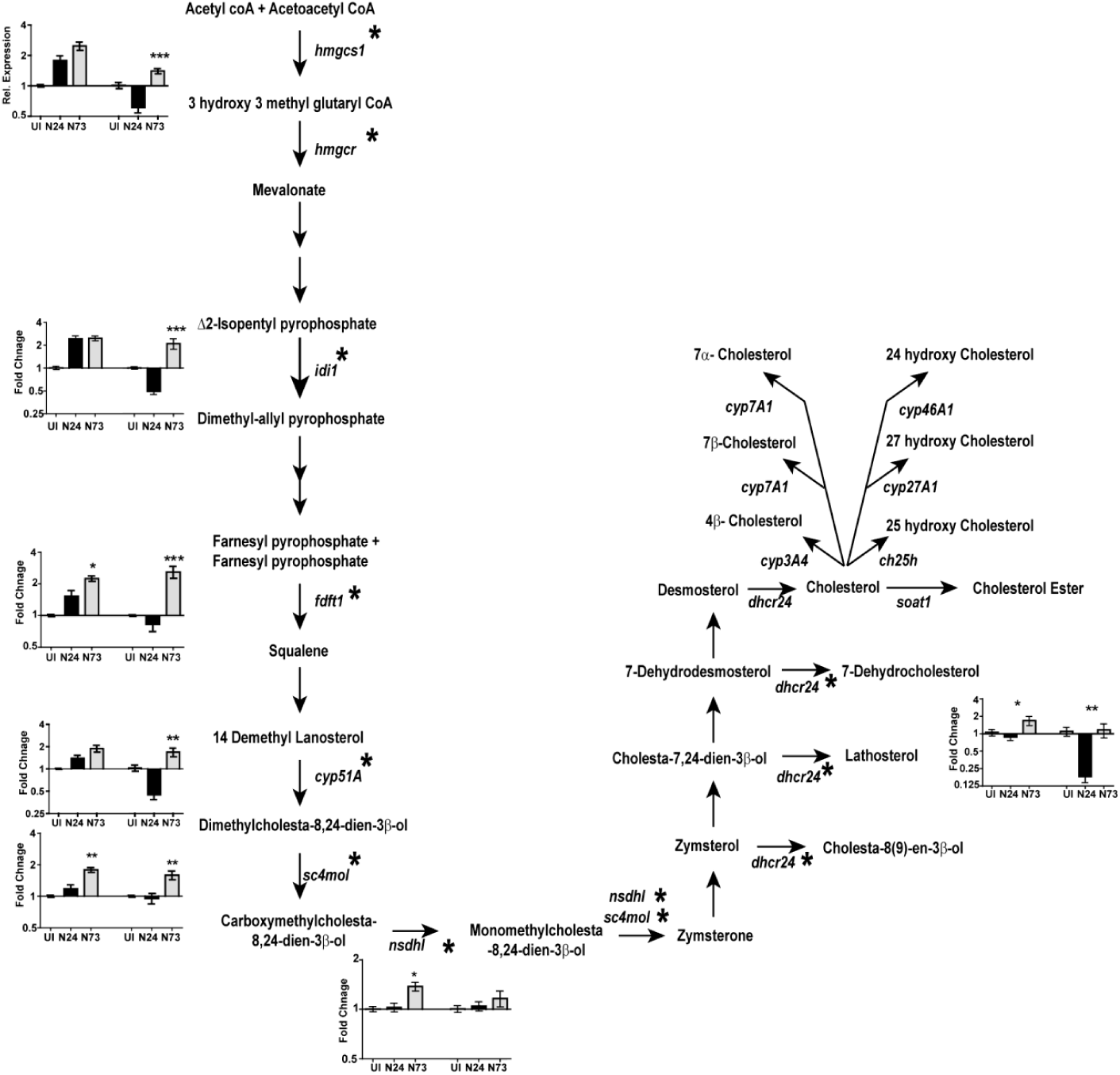
Expression of genes involved in cholesterol biosynthesis in macrophages infected with N24 or N73 at 6h and 24h (shaded) estimated by qRTPCR. The relative expression values ± SE (with respect to GAPDH) in triplicate assays of (N=2/3) is shown.

**Fig S3:**
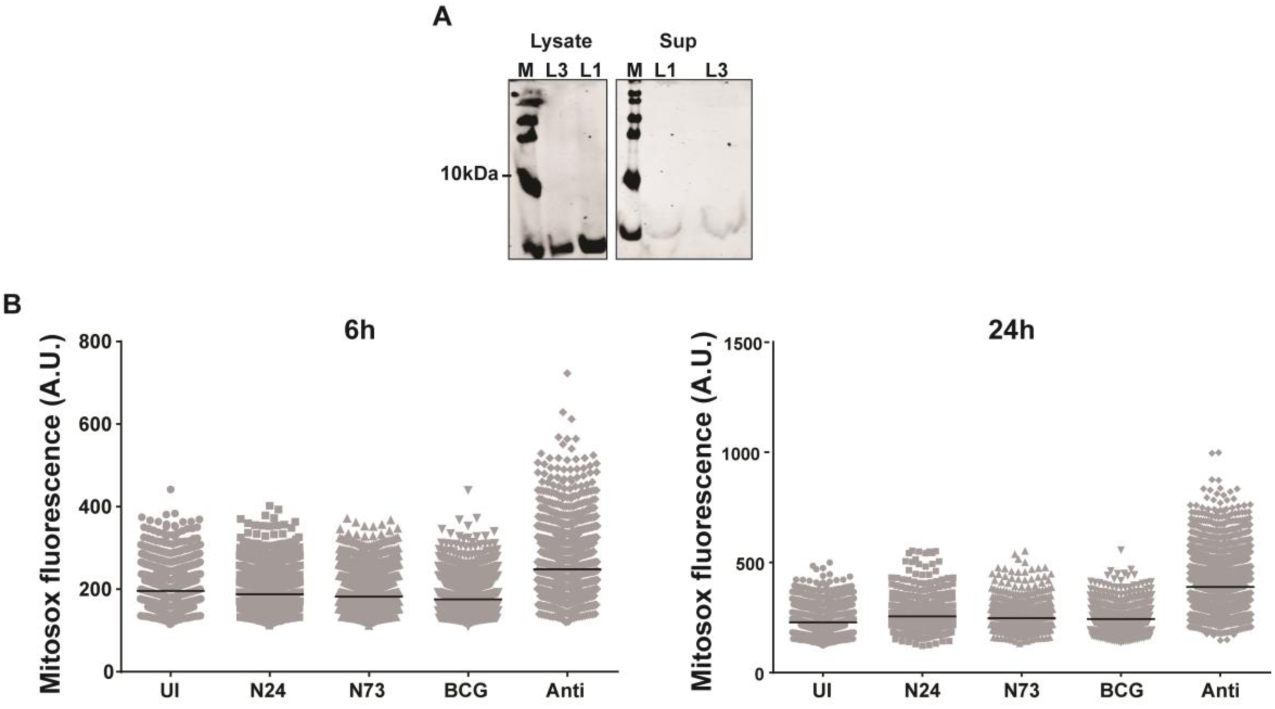
Analysis of factors responsible for type I IFN induction in macrophages following Mtb infection: A) ESAT6 expression in culture supernatant and lysates of Mtb N24 and N73. Expression was detected by immunoblotting with HYB76-8 monoclonal antibody. B) Quantitation of mitochondrial ROS by mitosox staining following infection with mycobacteria or treatment of Antimycin A (+ control) at 6h and 24h.

**Fig S4:**
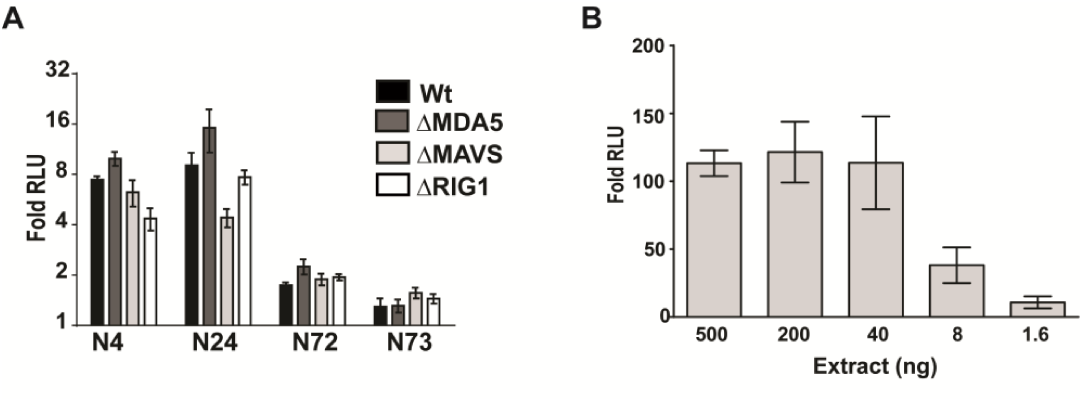
Mechanism of type I IFN induction in Mtb infected macrophages. A) Relative luminescence in RAW WT and KO macrophages infected with L3 and L1 Mtb at 24h post infection. B) Dose dependent IRF induction by Mtb crude extracts at 6h.

**Fig S5:**
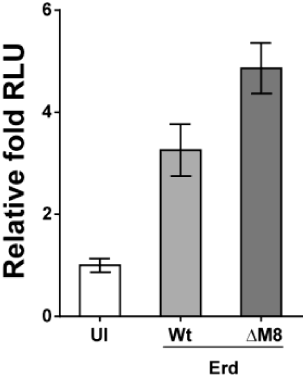
Evaluation of a putative role for Mtb sulfolipids in IFN induction. A) Relative luminescence in THP1 macrophages infected with Wt and MmpL8 mutant strains of Erdman at 24h p.i.

